# Site- and Structure-Specific Characterization of Glycoproteins of H11: Potent Vaccine Candidates against Parasitic Worm *Haemonchus*

**DOI:** 10.1101/2025.08.31.670727

**Authors:** Xin Liu, Feng Liu, Hui Liu, Lisha Ye, Yao Zhang, Wenjie Peng, Nishith Gupta, Min Hu, Chunqun Wang

**Author notes:** Corresponding authors: Min Hu, Email address, Chunqun Wang.

## Abstract

Parasitic worms (helminths) infections pose significant threats to global health and livestock economies. While mass spectrometry (MS)-based glycomics have revealed that helminths express complex, immunomodulatory N-glycans, site- and structure-specific characterization of intact glycopeptides remain challenging. Here we employed advanced MS-based intact glycoproteomics to explore the N-glycosylation profile of H11 antigen – an important vaccine antigen from pathogenic parasite *Haemonchus contortus*. We successfully identified seven glycosylated aminopeptidases carrying 19 N-glycosylation sites with 31 distinct N-glycan structures. Notably, 13 N-glycopeptides were significantly enriched by H11-induced protective IgG antibodies. Our findings revealed a high degree of structural heterogeneity and abundant core fucosylation among the identified N-glycopeptides. Additionally, molecular docking studies demonstrated those IgG-recognized N-glycopeptides are situated on the protein surface or adjacent to the substrate-access channels, strongly indicating their potential as antigenic epitopes. Overall, our work represents the first comprehensive glycoproteome of an economically important parasitic worm. These results hold important implications for the rational design of vaccines against *H. contortus* and other related metazoan parasites.

## 1 Introduction

Asparagine (N)-linked glycosylation of proteins is one of the most prevalent and functionally significant post-translational modifications (PTMs) in eukaryotic organisms. It plays essential roles in protein folding, molecular trafficking, receptor-ligand recognition, immune regulation, and disease progression (1, 2). This PTM exhibits substantial structural diversity, as glycans with various compositions can be covalently attached to multiple asparagine residues within polypeptides (3, 4). Mass spectrometry (MS) has emerged as the foundation of glycomics and glycoproteomics research. However, conventional MS-based approaches often rely on enzymatic deglycosylation. While this approach facilitates the analysis of glycan structures, it also erases valuable site-specific information regarding the glycoprotein heterogeneity (5, 6). Consequently, it is still challenging to determine whether certain glycosylation sites have a preferentially associate with specific glycan structures. In recent years, advanced MS-based intact glycoproteomic approaches have been developed to simultaneously preserve both the peptide backbone and the attached glycan moieties, enabling comprehensive mapping of site-specific glycosylation (7). Nevertheless, these powerful tools remain underutilized in many biological systems and organisms.

Parasitic worms (helminths) are a major global health and agricultural burden, causing chronic infections in humans and animals. Control strategies rely heavily on anthelmintic drugs, but widespread and often indiscriminate usage has led to increasing drug resistance, especially among gastrointestinal nematodes (8–10). Although substantial efforts have been made to develop vaccines against helminths, most candidates – whether native or recombinant – have failed to provide consistent and effective protection (11, 12). Notably, many studies suggest that helminth native glycoproteins can contribute significantly to host immunogenicity, with glycosylation patterns potentially influencing antigenicity and vaccine efficacy (13–15).

A key example is Barbervax®, the only commercially available gastrointestinal nematodes vaccine against *Haemonchus contortus* – a highly pathogenic gastric nematode in livestock, which has demonstrated high efficacy (75–95%) in field trials (16). The protective efficacy of this vaccine is attributed to the native intestinal membrane glycoproteins, predominantly composed of H11 aminopeptidase isoforms (17). However, attempts to produce effective recombinant H11 forms have been unsuccessful (18, 19), due to an incomplete understanding of the native protein glycosylation and immunodominant epitopes. Our recent findings revealed glycomics and glycoproteomic of H11, and demonstrating protective immunity elicited by H11 is closely associated with its N-linked glycan structures (15), However, the comprehensive linkage information between the diverse glycans and their attachment to multiple asparagine residues remains unknown.

Given that the structural heterogeneity of glycoproteins, this study employed advanced MS-based intact glycoproteomic techniques to perform the first comprehensive site-specific glycosylation analysis of H11 antigen. Our work highlights the significance of comprehensive glycoproteomic profiling in parasite biology and provides new insights for the rational design of vaccines targeting metazoan parasites.

## 2 Materials and methods

### 2.1 Biological reagents and chemicals

Con-A sepharose 4B was purchased from GE Healthcare (Pittsburgh, USA). Acetonitrile (ACN), dithiothreitol (DTT), trifluoroacetic acid (TFA), urea (57-13-6), Tris (2-carboxyethyl) phosphine (TCEP), iodoacetamide (IAA), ammonium bicarbonate (NH_4_HCO_3_) and trypsin NBr™ sepharose 4B were purchased from Sigma Aldrich (St. Louis, MO, USA). Ammonium bicarbonate (NH_4_HCO_3_) and bicinchoninic acid (BCA) protein assay kit were purchased from Sangon Biotech (Shanghai, China). Ultrapure water was produced on site by Millipore Simplicity System (Billerica, MA, USA).

### 2.2 Sample preparation of H11 antigen

Native H11 antigen was isolated from adult *H. contortus* using ConA-sepharose as previously described (20). Briefly, worms were homogenized in ice-cold phosphate-buffered saline (PBS, pH 7.4) for 25 min using a glass homogenizer. The homogenate was centrifuged at 12,000 g for 25 min. The pellets were then extracted four times with 1% (v/v) Thesit in PBS and filtered (0.45 µM). The extract was loaded on a ConA-Sepharose column. The column was washed 3 times (20 mM Tris-HCl), followed by elution with 200 mM of methyl α-D-mannopyranoside and methyl α-D-glucopyranoside solution. The fractions containing H11 were pooled and filtered (0.22 µM). This final filtrate was designated the native H11 antigen. The protein purity was assessed by Coomassie blue staining on SDS-PAGE and the concentration was determined using a BCA kit.

### 2.3 Anti-H11 specific IgG antibodies

The IgG antibodies used in this study were derived from our previously established methodology (15). Briefly, IgG were purified by Protein A+G agarose from sera of goats immunized with untreated natural H11 antigen (NA group) or deglycosylated H11 antigen treated by sodium periodate to destroy glycans (PI group).

### 2.4 Sample preparation of intact N-glycopeptides

The workflow of analysis of intact N-glycopeptide is shown in Fig 1, including sample preparation and MS analysis. In brief, ConA-purified H11 glycoprotein was reduced with 10 mM DTT at 55 °C for 30 min, alkylated with 20 mM IAA at RT in the dark for 20 min, and quenched by an additional 10 mM DTT. The protein was then dissolved in 5 mL 50 mM −H_4_HCO_3_ and digested with 20 μg trypsin at 37 °C for overnight. A homemade C18 Solid-Phase Extraction (SPE) tip (Jupiter C18, particle size 15 μm, pore size 300 Å) was used for desalting. The C18 tip was pretreated sequentially by 80% ACN with 5% TFA and then by 0.1% TFA. The digestion solution was loaded and washed with 0.1% TFA aqueous, and peptides were eluted by 100 μL 50% ACN with 0.1% TFA and then 100 μL 80% ACN with 5% TFA. The eluted peptides were dried in a SpeedVac and resuspended in 100 μL 80% ACN with 5% TFA. Intact N-glycopeptides were enriched with homemade SPE columns packed with ZIC-HILIC particles (Merk Millipore, particle size 5 μm, pore size 200 Å). The column was first equilibrated with 0.1% TFA aqueous and then followed by 80% ACN with 5% TFA. The peptide mixtures were loaded on the ZIC-HILIC SPE column. After washing by 80% ACN with 5% TFA (21), the peptides were eluted with 100 μL of 0.1% TFA aqueous and with 50 μL of 50 mM NH_4_HCO_3_. The eluates were combined, dried in a SpeedVac, and resuspended in 40 μL of H_2_O for downstream LC-MS/MS analysis.

**Figure 1.**
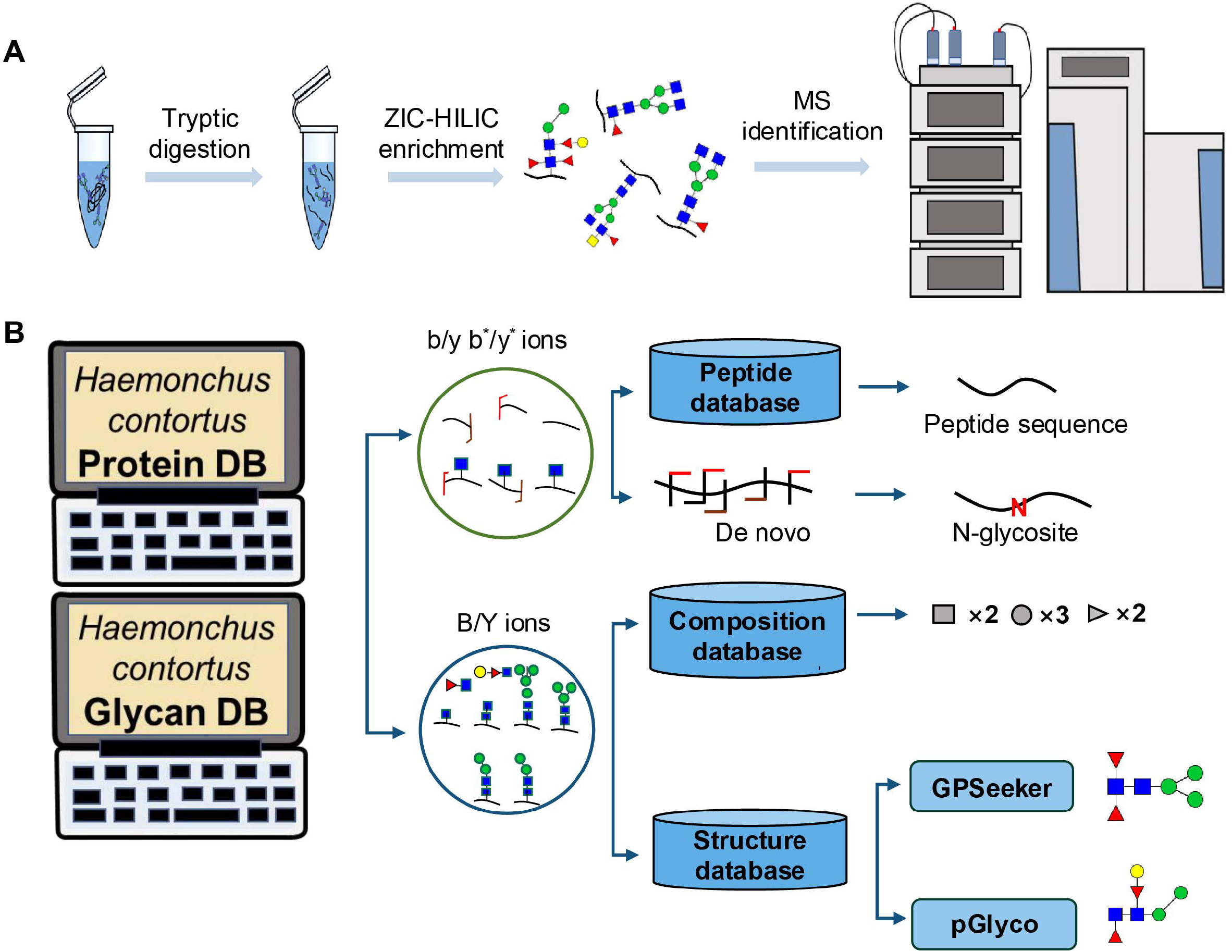
Workflow of mass spectrometry (MS)-based intact glycoproteome analysis of H11 antigen. (A) H11 glycoproteins are performed trypsin digestion, ZIC-HILIC enrichment and MS identification. (B) Comprehensive N-glycoproteome databases based on known H11 glycome and protein isoforms. Data analysis is employed b/y product ion to determine peptide backbone from the GlcNAc-containing site, and performed GPSeeker and pGlyco to map the structure-diagnostic B/Y product ions from the N-glycan moieties.

### 2.5 Nano-RPLC ESI-MS/MS analysis

Samples were analysed on an Ultimate 3000 RSLC nano HPLC system (Dionex, Sunnyvale, CA, USA) coupled to a Q Exactive Orbitrap mass spectrometer (Thermo Scientific) operating in the positive ion mode. Chromatographic separation was performed on a C18 (ZORBAX 300SB, 5 μm, 300 Å; Agilent Technologies, Santa Clara, CA, USA) analytical column (60 cm x 360 μm o.d. × 75 μm i.d.) at a flow rate of 300 nL/min with the mobile phases of 99.9% H_2_O with 0.1% FA (A), 99.9% ACN and 0.1% FA (B). The LC gradient was set as follows: 2% B, 12 min; 2%-40% B, 180 min; 40%-95% B, 5 min; 95% B, 5 min; 95%-2% B, 5 min; 2% B, 20 min.

The ESI conditions were set as follows: spray voltage 1.9 kV and capillary temperature 300 °C. Full-scan mass spectra were acquired in the range of 700–2000 m/z with major ESI source settings: mass resolution 60 k, automatic gain control (AGC) target 3×10^6^, max ion injection time 20 ms; MS/MS spectra were acquired on Top20 data-dependent acquisition mode with the following settings: mass resolution 30 k, AGC target 5×10^5^, max ion injection time 250 ms, isolation window 3 m/z, higher energy collisional dissociation (HCD) with stepped normalized collision energies 20%, 30% and 40%; dynamic exclusion 20 s.

### 2.6 Database search and identification of intact N-glycopeptides

All intact N-glycopeptides deposited in the database were constructed *in silico* by exhaustively combining the computed N-glycosite-containing trypsin-digested peptides with a total of 50 PNGase F-released *H. contortus* N-glycan structures (Supplementary Table 1) (15) Computed N-glycosite-containing peptides were generated utilizing proteins from Uniprot (https://www.uniprot.org/) with the filtration of organism name *H. contortus* aminopeptidase and a total of 13 reported proteins were finally selected (Supplementary Table 2) (22). The intact N-glycopeptide database was divided into two discrete subsets to enable parallel interrogation through two specialized search engines, with GPSeeker and pGlyco 3.0 being respectively employed for independent analyses.

Thus, each acquired raw spectrum dataset was analyzed independetlyby engine GPSeeker and pGlyco 3.0 to generate two sets of intact N-glycopeptide identifications (IDs) with false discovery rate (FDR) < 0.01. Each set of IDs was then processed by removing repetitively reported N-glycopeptide IDs and filtering IDs until no MS2 spectrum was assigned to multiple N-glycopeptide matches in each engine report. Finally, remaining IDs from two engines were merged to obtain an engine-union qualitative IDs list. It is worth noting that a little fraction of matched MS2 spectra were annotated to different N-glycopeptides independently by these two engines due to the formula isomers between the split databases. To ensure the unique N-glycopeptide match of each MS2 spectrum, the ID with more matched peptide and glycan fragments was kept as the true interpretation.

### 2.7 Silver staining analysis of IgG-recognized H11 proteins

Anti-NA, anti-DN or anti-PI IgG antibody (50 μg/mL, 1 mL) was incubated with Protein A+G beads at room temperature for 30 min. Subsequently, native H11 samples were added to above mixture and further incubated at 4 °C for 3 h. The magnetic beads were then separated with a magnet followed by washing with TBS. Finally, the magnetic beads were immersed in SDS-PAGE sample loading buffer and boiled for 10 min. Following magnetic separation for 10 s, the supernatant was collected for silver staining.

### 2.8 Identification of IgG-bound glycopeptides

The workflow was carried out according to a modified protocol (23). Briefly, 250 μg of anti-NA or PI IgG were dissolved in 100 mM NaHCO_3_ buffer (pH 8.5) containing 500 mM NaCl, respectively. Meanwhile, 0.5 mL of CN-Br Sepharose beads were swollen in 1 mM HCl at 4 °C for 30 min. It was washed with 10 mL ddH_2_O to remove the HCl, and followed by 1 mL NaCl/NaHCO_3_ buffer (pH 8.5) to activate the resin and immediately transferred to the ligand solution in the coupled buffer above each IgG solution. After incubation of IgG antibodies and agarose beads at room temperature for 2 h, the mixture was centrifuged at 13,000 g for 1 min, and the supernatant was discarded. The agarose beads were then rinsed with 2 mL NaHCO_3_/NaCl buffer to remove the unbound antibodies. Finally, we harvested the supernatant which did not bind to the anti-PI IgG antibody, and the supernatant was bound with anti-NA IgG at room temperature for 4 h. After centrifuging, the pellets were thoroughly cleaned with 25 mM ammonium acetate (pH 7.5) for 4 times to remove unbound proteins. Finally, specifically NA-IgG bound proteins were eluted by incubation with 1 ml of 25 mM (pH 3.0) ammonium acetate eluent for 30 min. Finally, This MS-based intact glycopeptides analysis was conducted based on above established protocol (see sample preparation of intact N-glycopeptides 2.4 and Nano-RPLC ESI-MS/MS analysis 2.5).

### 2.9 Analysis of intact N-glycopeptides

Quantitative analysis of each N-glycopeptide was calculated as the sum of the intensities of peaks in the isotopic fingerprinting peaks of the corresponding intact N-glycopeptide precursors regardless of the applied search engine. IDs with a lack of the precursor top peak matches were eliminated from the processes of quantitative analysis. The relative abundance of each quantitative N-glycopeptide ID was carried out using diverse normalization methods including sum normalization, max-min normalization, mean normalization, Z-score normalization, and log normalization. The quantitative IDs list was constructed by selecting the corporately identified N-glycopeptides with the tolerance of missing identification in at most 50% of the samples in the same group, and missing values were imputed by its neighboring value multiplied by a normalization factor. The normalization factor was calculated by first filtering out a serial of inter-sample shared N-glycopeptide with extremely high quantitative linear correlation and then computing the ratio between samples as the normalization facto.

### 2.10 Molecular docking

The tertiary structure of the H11 protein PDB files were downloaded from the AlphaFold Protein Structure Database (https://alphafold.ebi.ac.uk/); SDF files of the small molecule Leu-pNA (CNS No. 4178-93-2) were downloaded from the PubChem database and converted to PDB files using PyMOL. After hydrogenation, charge calculation, merging non-polar hydrogen and other operations, generated structure was saved as pdbqt file. Run the algorithm to select the local search parameters to calculate more than 50 times. Sorting by energy, select the compound with the lowest binding energy in the docking results. Autodock programme was used for docking, while PyMOL was used to visualize the docking data and the amino acid residues that form hydrogen bonds in the 4 Å range. Surface position distribution was analyzed using the NetSurfP programme (https://services.healthtech.dtu.dk/services/NetSurfP-3.0/). N-glycopeptide enrichment analyses were performed using the Hiplot (https://hiplot.com.cn/home/index.html/) and Microbiotics websites (http://www.bioinformatics.com.cn/).

## 3 Results

### 3.1 Experimental Design

We have previously analysed the N-glycome and N-glycoproteome profile of *H. contortus* (15, 24), but the precise correlation between specific glycosylation sites and their corresponding glycan structures remains unresolved. To address this gap, we characterized comprehensive site- and structure-specific N-glycosylation in H11 components. We implemented an integrated N-glycoproteomics workflow combining tryptic digestion, ZIC-HILIC enrichment and MS identification (Figure 1 and Supplementary Tables 1, 2). In the analysis of MS spectra, we employed b/y ions to identify the GlcNAc-carrying peptide fragments, and GPSeeker and pGlyco 3.0 algorithms were applied to characterize the glycan compositions via B/Y ion mapping (Figure 1B), which has been well established in N-glycoproteome analysis (25, 26).

### 3.2 Identification of intact N-glycopeptides of H11 antigen

We identified 202 intact N-glycopeptides mapping to 19 N-glycosylation sites (Figure 2A and Supplementary Table 3). The conserved N-x-T/S sequons (x ≠proline) demonstrated a marked preference for threonine residues over serine (Figure 2B), consistent with our previous findings in *H. contortus* (15, 24). Isoform-specific analysis showed that H11 antigen contained five known isoforms (H11, H11-1, H11-2, H11-4 and H11-5) (12, 27–29) and two aminopeptidase molecules (AP-3 and AP-6) (22), with H11, H11-1, H11-4, H11-5 and AP-6 showing predominant expression (Figure 2C).

**Figure 2.**
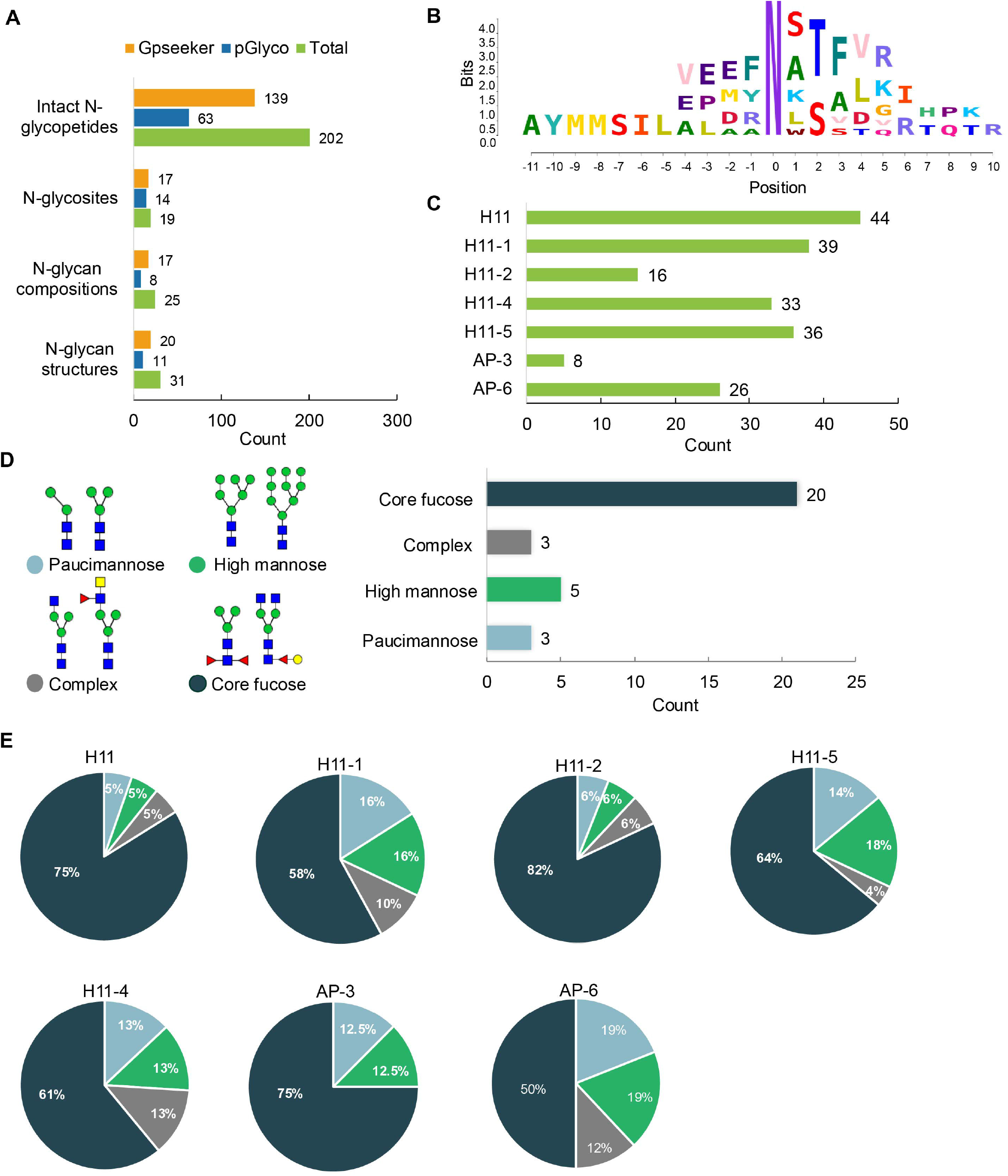
Characteristics of the intact N-glycopeptide profile of H11 antigen. (A) Information on the numbers of intact N-glycopepetides, N-glycosites, N-glycan compositions and N-glycan structures from GPSeeker and pGlyco techniques, respectively. Data are derived from two independent experiments. (B) Motif analysis of all identified N-glycopeptides. (C) The relative percentage of intact N-glycopeptides identified in each H11 aminopeptidase. (D) The representative glycan structures and the percentage distribution of N-glycan structure classes. (E) The percentage of four types of N-glycans identified in each H11 aminopeptidase.

Glycomic profiling revealed 25 distinct N-glycan compositions corresponding to 31 structures (Figure 2A and Supplementary Table 3), categorized into four types, i.e., paucimannose (Hex_2-4_HexNAc_2_), high-mannose (Hex_5-9_HexNAc_2_), complex (Hex_3-4_HexNAc_3-5_Fuc_0-2_) with or without core fucose (Hex_2-4_HexNAc_2_Fuc_0-3_) (Figure 2D). Notably, 65% (n=20) of identified structures exhibit α1,3- and/or α1,6-fucosylation (Figure 2D and Supplementary Table 3). In H11, H11-2 and AP-3, over 75% structures are core fucosylated (Figure 2E). The core fucosylation has been demonstrated to function as a pathogenic and immunological glyco-determinant for helminth N-glycans (15). Hence, our intact N-glycopeptide identification deepens the understanding that ‘non-mammalian’ glycans, including α1,3-core fucose, could be critical elements for H11 antigen to stimulate the host’s immune response.

### 3.3 Heterogeneity analysis of H11 glycoproteins

Due to the non-template driven nature of glycosylation *in vivo*, natural proteins are normally glycosylated with diverse glycan structures, referring to macro-heterogeneity (i.e., more than one glycosite in one protein) and micro-heterogeneity (i.e., more than one glycan structure at individual glycosite) (30). Our heterogeneity analysis of H11 glycoproteins revealed 19 N-glycosylation sites hosting 31 distinct glycoforms. Around 89% of glycosylation sites exhibit micro-heterogeneity (Figure 2A). We observed that there are three hypervariable sites (H11 N^227^, H11-1 N^235^ and H11-4 N^226^), each of them bearing 16 glycoforms (Figure 3A). According to macro-heterogeneity, five glycoproteins (i.e., H11, H11-1, H11-4, H11-5 and AP-6) have more than three N-glycosites and the other two glycoproteins (i.e., H11-2 and AP-3) possessed only one glycosite (Figure 3B), which expands our previous reports on the glycosylation sites on six aminopeptidases with 13 glycosylation sites (15), suggesting that H11 antigen exists in a higher degree of heterogeneity.

**Figure 3.**
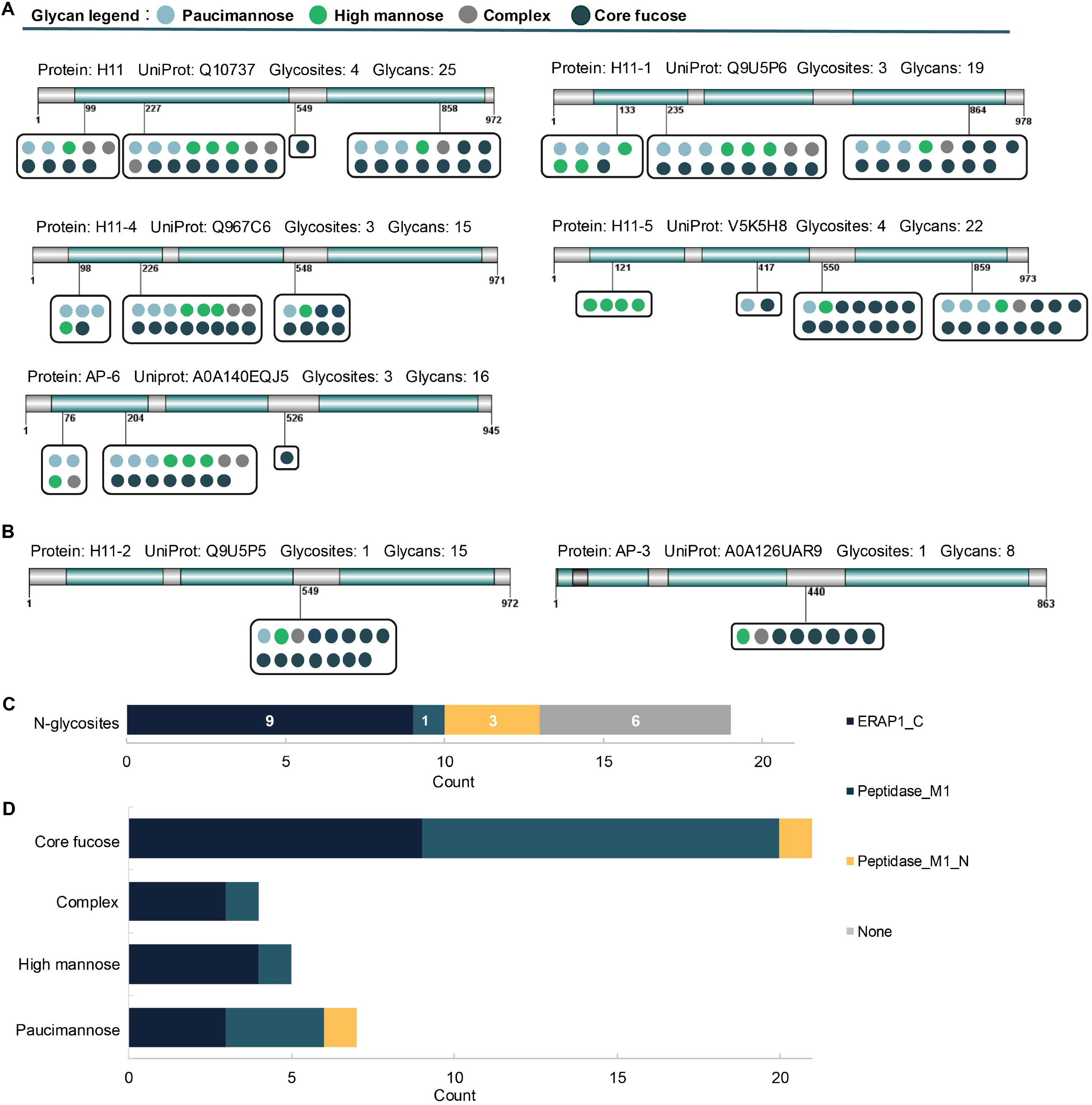
Heterogeneity and protein domain analysis of H11 glycoproteins. (A and B) H11 aminopeptidase with more than two or only one glycosylation sites, showing protein accession number in UniProt, number of glycosylation sites and glycan structures. (C) The number of N-glycosylation sites localized in each domain. (D) The number of different glycan types in each protein domain.

To assess the structural features of glycosylation site on H11 molecules, we further analysed protein domain based on the UniProt database. Our data revealed that of which 13 glycosylation sites were in three main structure domains, including the ERAP1_C domain, Peptidase_M1 domain, and peptidase_M1_N domain (Figure 3C). Around 47% (*n* = 9) of glycosylation sites were in the ERAP1_C domain, and three sites were in the peptidase_M1_N domain, as well as only one in the Peptidase_M1 domain (Figure 3C). ERAP1_C has been reported to be a concave surface toward the aminopeptidase activity, and that the arginine-aspartate motif in the ERAP1_C domain has been implicated as critical for substrate targeting (31, 32). Compared with the glycan motifs between the different domains, we observed that a high proportion of fucosylation occur in the ERAP1_C and Peptidase_M1 (Figure 3D). Althought currently there is no functional study of protein fucosylation associated with aminopeptidase activity, it is worth to be investigated further.

### 3.4 Identification of potential B-cell epitopes within H11 glycopeptide

Given that the importance of N-glycopeptide epitopes, we further explored potential B-cell epitopes by H11-induced IgG antibodies. Our silver staining analysis clearly demonstrated glycosylated H11 was strongly recognized by protective NA-group IgG, but not by PI-group IgG (Figure 4B). Furthermore, in combination with immunoprecipitation study and glycoproteomic analysis (Figure 4A), we identified 317 intact N-glycopeptides (Figure 4C and Supplementary Table 4). Among these, 13 glycopeptides exhibited significant enrichment by protective IgG (Figure 4D and Supplementary Table 5). These glycopeptides are distributed across seven glycosylated aminopeptidases and are associated with nine distinct N-glycan motifs, all featuring with core fucosylation. Such motifs include difucosylated form (α1,3- and α1,6-linked), galactosylated α1,6-core fucose and relatively uncommon trifucosylated motifs (Table 1). Notably, we observed VEEF<u>N</u>ATALK peptide sequence bore three distinct N-glycoforms (Table 1). Meanwhile, our findings revealed that N-glycan chains (Hex_4_HexNAc_4_Fuc_1_) were attached to three peptides (A<u>N</u>WTVTVIHPK, <u>N</u>LTFDGR and VEEF<u>N</u>ATALK), while (Hex_2_HexNAc_2_Fuc_2_) were associated with AFNATSLQITQTR ALDRNSSFVR VEEFNATALK (Figure 5A and Table 1). These results underscore the remarkable heterogeneity of those potential glycopeptide epitopes present within the H11 antigen.

**Figure 4.**
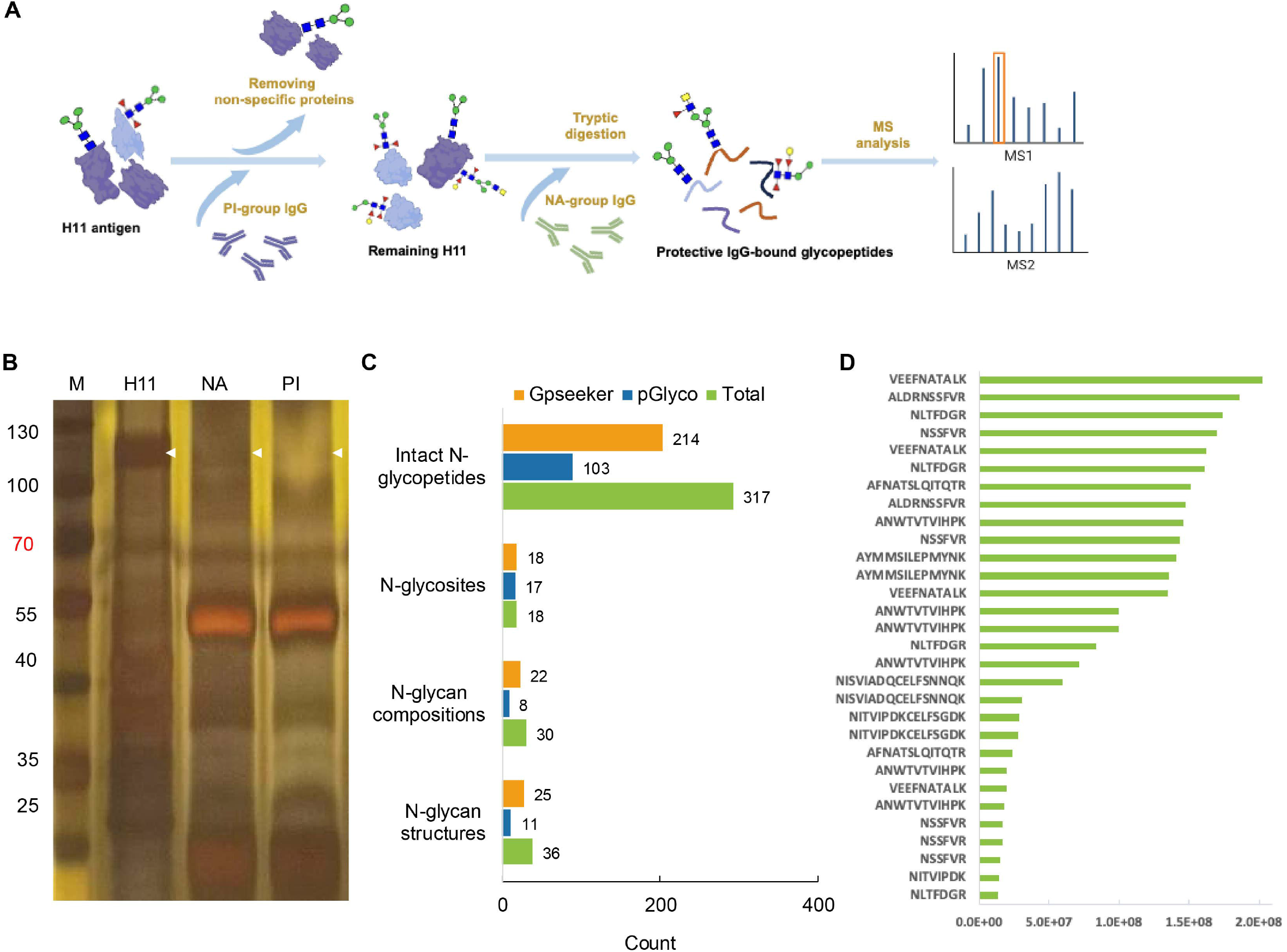
Identification of potential epitopes on H11 glycopeptide. (A) Schematic representation of the experimental procedure of identification of IgG-bound glycopeptides. (B) Silver staining analysis of native H11 antigen and NA or PI-group IgG-bound H11 antigen. NA-group IgG antibody is induced by untreated natural H11 antigen, and PI-group IgG antibody is made by deglycosylated H11 antigen without glycans modification. (C) Information on the numbers of intact N-glycopepetides, N-glycosites, N-glycan compositions and N-glycan structures from identified IgG-bound glycopeptides. Data are derived from two independent experiments. (D) Enrichment analysis of identified IgG-bound glycopeptides (Only showing 30 obvious enriched glycopeptides).

**Figure 5.**
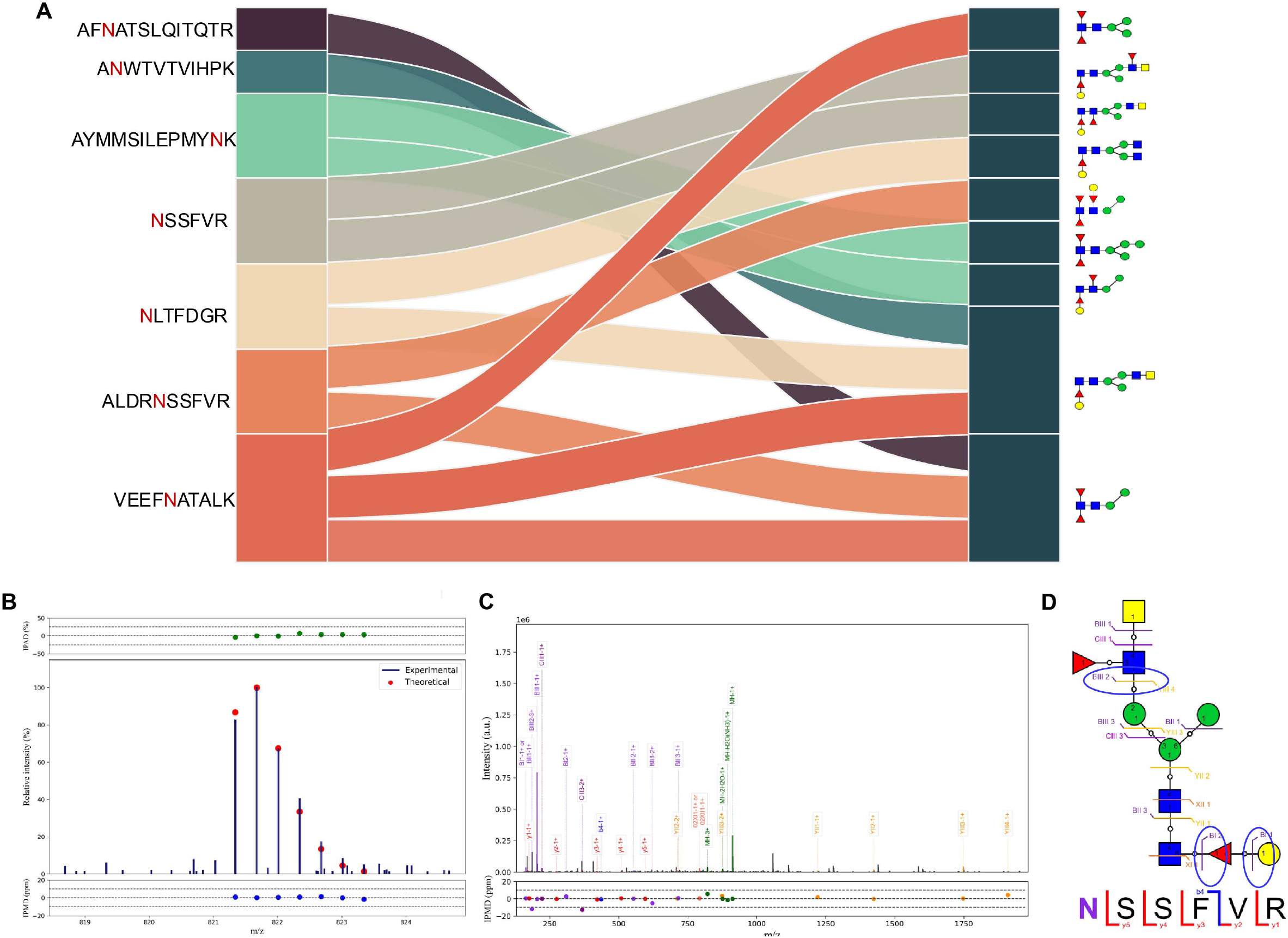
Analysis of potential epitopes on H11 glycopeptide. (A) correlation analysis between identified glycopeptide and their linked N-glycan structures. (B) The isotopic envelope finger printing maps of the N-glycopeptide NSSFVR-01Y(61F41L)41Y41M(31M21Y(31F)41V)61M precursor ions. (C) Annotated MS/MS spectrum of matched fragment ions. (D) Graphical fragmentations of the peptide backbone and the N-glycan structure isomers.

Furthermore, we identified a previously unreported glycan (Hex_4_HexNAc_4_Fuc_2_) in native H11 sample. MS analysis showed that BIII2 ions was corresponding to a glycan motif composed of two HexNAc residues and one Fucose, while BI2 and BI1 fragment ions were indicative of a hexose attached to a core fucose (Figures 5B, C). Database analysis suggested that this glycan consists of a galactosylated α1,6-linked fucose motif with a LDNF unit (Figure 5D). Its absence in prior H11 antigen analyses suggest that its detection here may result from selective enrichment by protective IgG antibodies. This evidence suggests that N-glycopeptide epitopes have the potential to sever as valuable candidates for the development of vaccines against *H. contortus* and related parasitic worms in the future.

### 3.5 Molecular docking and visual analysis of potential epitopes in H11 glycopeptide

Protective antibodies can inhibit enzyme activity through steric hindrance and conformational changes, depending on the spatial relationship between their target epitopes and the enzyme’s active site (33). In this study, we employed Autodock and PyMOL to predict the substrate-binding regions in all identified H11 glycoproteins. Subsequently, through 3D structure visualization, we analyzed the position interactions between antibody and its binding epitope at the atomic level. Molecular docking revealed that the substrate-binding interface is situated in the protein’s internal cavity, and the substrate forms multiple hydrogen bonds with key amino acid residues of various H11 isoforms within a 4 Å radius, such as Glu402 and Tyr466 in a H11 protein (Figures 6A, B). Structural visualization allowed us to explore the spatial arrangement between the aminopeptidase active site and antibody-binding epitopes (Figures 6C, D and Supplementary Figure 1). In the H11 protein, antibody-binding sites surround the substrate-binding channel (Figure. 6C). In contrast, in the H11-2 protein, the antibody-binding sites are distal to the active center (Figure 6D). Based on these findings, we propose that protective IgG may target those glycopeptide epitopes adjacent to the substrate-binding region, inducing steric blockade or allosteric modulation that disrupts substrate hydrolysis. This mechanism could explain how antibody-mediated interference with aminopeptidase activity contributes to immune protection. Further research on these glycopeptide epitopes will provide critical insights for designing anthelmintic drugs or vaccines against *H. contortus* or related parasitic worms.

**Figure 6.**
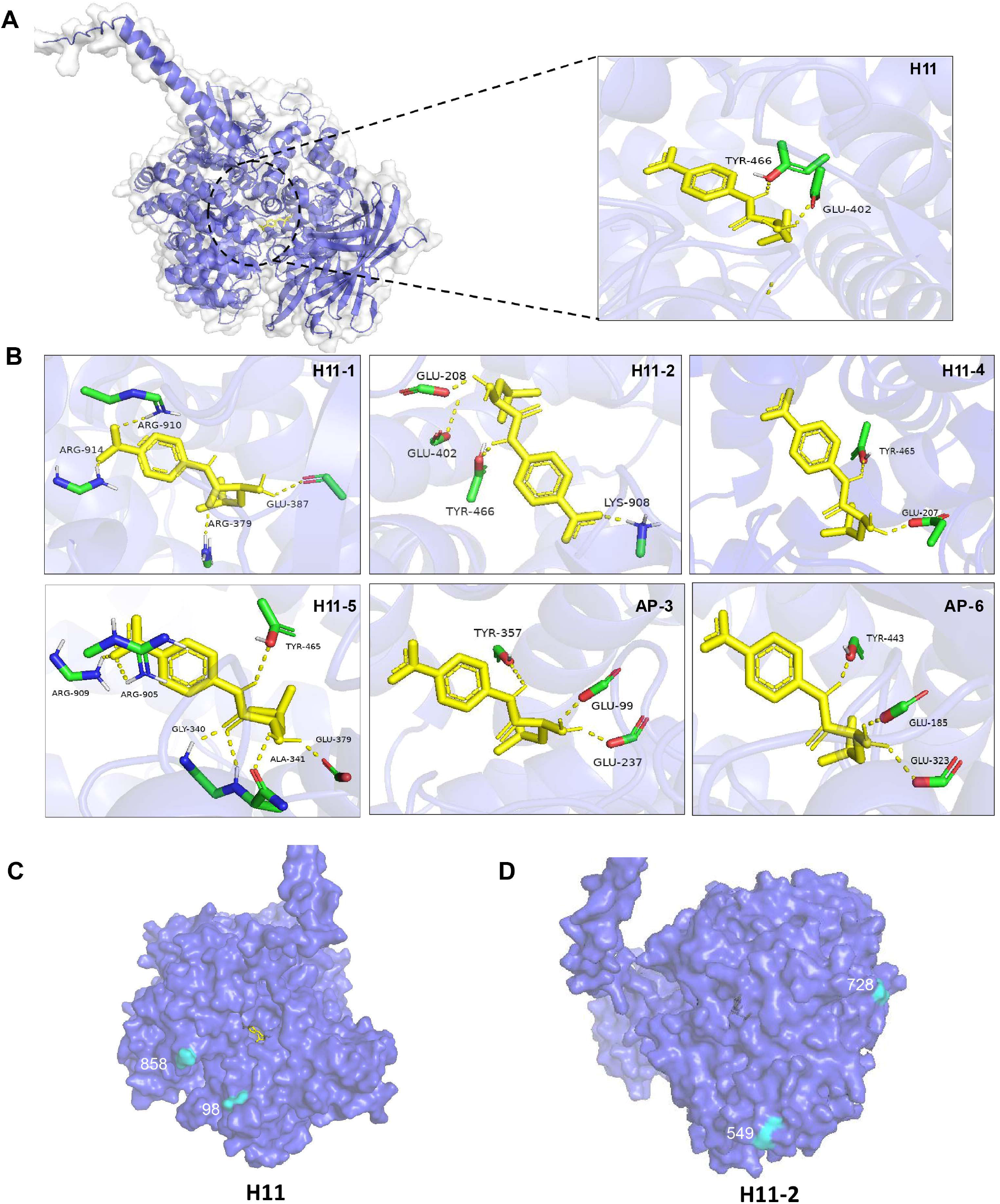
Molecular docking and visualization of identified H11 molecules. (A) Complex structure of H11 subtype and substrate (Leu-pNA). (B) Local diagram of H11 aminopeptidase amino acid residues interacting with substrate binding. (C and D) 3D structures of H11 and H11-2 are shown in Cartoon styles.

## 4 Discussion

H11 antigen, a well-characterized immunoprotective molecule from a parasitic nematode. In its native form, H11 consistently confers high levels of protection (usually 75% to 95%) in immunized animals (17). Although substantial academic–industry collaborations, recombinant H11 proteins produced in both prokaryotic and eukaryotic systems failed to elicit comparable levels of protective immunity in animal models (22, 34). To elucidate the mechanisms underpinning its potent immunogenicity, our previous work defined the glycome and glycoproteome of the native H11 antigen and highlighted the critical role of glycan structures (15). However, detailed site- and structure-specific profile of glycoproteins of H11 remains unknown.

Here we performed the first comprehensive N-glycopeptides analysis of *H. contortus* H11 antigen. Using MS-based intact glycoproteomics, integrating GPSeeker and pGlyco platforms, our analysis identified seven aminopeptidases including five H11 isoforms (H11, H11-1, H11-2, H11-4 and H11-5) and two additional aminopeptidase molecules (AP-3 and AP-6). These molecules collectively harbored 19 glycosylation sites bearing 31 glycan structures, showing that H11 glycoproteins exists in a higher degree of heterogeneity. So far, decoding the intact glycoproteome of parasitic worms remains a significant technical challenge, due to the complex and species-specific nature of their glycans. Helminth glycans often feature highly branched architectures, non-mammalian residues, and extensive fucosylation (35, 36), which impede standard analytical workflows. Despite these challenges, analysis of glycosylation at the site-specific level is still essential, as intact glycoproteins can profoundly influence parasite biology, immune recognition, and vaccine efficacy (37, 38).

We demonstrated that 13 N-glycopeptides were enriched by specific protective IgG antibodies. Upon further characterization, a high degree of structural heterogeneity and core fucosylation were revealed, including difucosylated, galactosylated α1,6-linked monofucosylated and uncommon trifucosylated motifs. Importantly, core fucose motifs have been implicated in the immunomodulatory activity of helminth glycoproteins. For instance, α1,3-core fucose has been specifically detected on glycoprotein omega-1 from *Schistosoma mansoni* eggs, which are associated with its Th2-inducing property (39). In addition, α1,3-core fucose on *H. contortus* excretory/secretory glycoproteins functions as B-cell epitope, as evidenced by the recognition of IgG antibodies from vaccinated animal serum (40). Previous studies have shown that the LDNF epitopes are targeted by IgG antibody in sheep vaccinated with *H. contortus* excretory/secretory glycoproteins (41). Similar motifs have been exploited as immunogenic glycans in the diagnosis and development of glycoconjugate vaccine for *S. mansoni* (42, 43).

Molecular docking studies demonstrated that these IgG-recognized N-glycopeptides are situated on the protein surface or adjacent to the substrate entry channels leading to the active site of the aminopeptidases. This spatial localization suggests a potential immunological mechanism whereby antibody binding may sterically hinder substrate access or enzyme activity, thereby disrupting key physiological functions of the parasite aminopeptidases. Our previous work has revealed this potential mechanism that protective IgG antibodies target glycans of aminopeptidases and inhibit the enzyme activity, which correlates with immunoprotection (15). Such interference could contribute to the protective efficacy observed in immunized animals. Moreover, the surface exposure of these glycopeptide epitopes enhances their accessibility to host immune surveillance, enhancing their relevance as immunodominant targets, although accurate modeling of glycopeptide epitopes with defined glycan modifications remains technically challenging. These findings support the hypothesis that glycopeptide epitope positioning are critical determinants of protective immunity, and further underscore the importance of preserving glycosylation patterns in the development of effective subunit vaccines. Although the use of glycopeptide epitopes for vaccine development has yet to be established in the field of parasitic worm, substantial studies have shown that N-glycopeptide can possess an effective immunogenicity (44). For example, HIV-neutralising antibodies targeting conserved gp120 glycopeptides have successfully elicited robust immune responses in animal models (45, 46). Thus, a comprehensive understanding of these H11 glycoproteins can therefore provide new molecular insights into protective immunity and guide the rational design of effective glyco-engineered subunit vaccines in the future.

In conclusion, the study conducted the first comprehensive N-glycopeptide analysis of native H11 using MS-based intact glycoproteomics. We identified seven glycosylated aminopeptidases with 19 glycosylation sites carrying 31 structurally diverse glycans. Thirteen N-glycopeptides were selectively enriched by protective IgG antibodies, revealing core fucosylated and structurally heterogeneous motifs, including immunologically relevant α1,3-core fucose. Docking studies showed these epitopes are surface-exposed or near active sites, suggesting antibody binding could block enzymatic function, contributing to protective immunity. These findings highlight the critical role of native glycosylation and epitope positioning in vaccine efficacy, supporting the potential of glyco-engineered subunit vaccines based on intact glycopeptide antigens. Further studies will fucose on synthesis of those immunodominant glycopeptide epitopes to evaluate their protective efficacy in animal models, paying the way for novel antihelmintic vaccine development.

## Data availability statement

All data generated or analysed during this study are included in this published article and its supplementary information files.

## Ethics statement

This study was reviewed and approved by the Animals Ethics Committee of Huazhong Agricultural University (permit HAZUGO-2025-0004).

## Authors’ contributions

**Xin Liu:** Writing – review & editing, Writing – original draft, Visualization, Validation, Software, Methodology, Investigation, Data curation. **Feng Liu:** Writing – review & editing. **Hui Liu:** Methodology, **Lisha Ye:** Methodology. **Yao Zhang:** Methodology, **Wenjie Peng:** Methodology, Writing – review & editing, Supervision. **Nishith Gupta:** Writing – review & editing, Supervision. **Min Hu:** Writing – review & editing, Supervision, Resources, Project administration, Funding acquisition. **Chunqun Wang:** Writing – review & editing, Writing – original draft, Visualization, Validation, Software, Methodology, Supervision, Resources, Project administration, Funding acquisition. All authors read and approved the final manuscript.

## Acknowledgments

We thank the Prof. Zhixin Tian from Tongji University for their assistance with intact glycopeptide analyses and technical support.

---

**Table 1.**
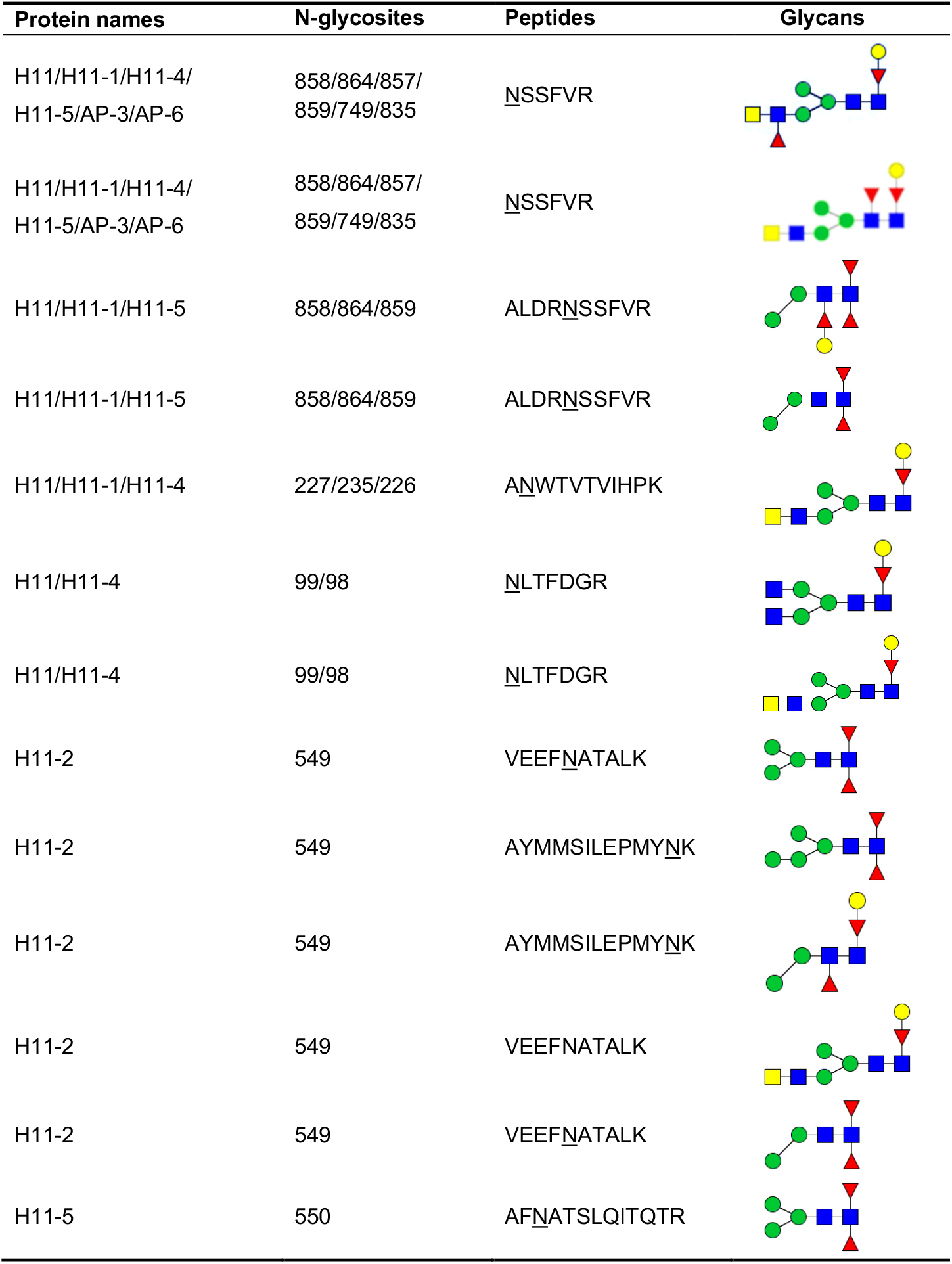
Mapping of protective IgG-bound intact N-glycopeptides of H11 antigen.

## Supplementary Information

**Supplementary Figure 1**. Molecular docking and visualization of identified H11 molecules. 3D structures (A-E) of H11-1, H11-4, H11-5, AP-3, and AP-6 are shown in Cartoon styles.

**Supplementary table 1**. Database of glycan structures and assignment to target GPSeeker and pGlyco search engines

**Supplementary table 2**. Aminopeptidases database based on previous known 13 aminopeptidase H11 molecules

**Supplementary table 3**. Information of intact N-glycopeptides identified from native H11 antigen

**Supplementary table 4**. Information of intact N-glycopeptides identified from IgG-bound H11 antigen

**Supplementary table 5**. Enrichment of intact N-glycopeptides identified from IgG-bound H11 antigen based on intensity information

