## Supplementary figures and images for "Site- and Structure-Specific Characterization of Glycoproteins of H11: Potent Vaccine Candidates against Parasitic Worm *Haemonchus*"

### Supplementary Figure 1

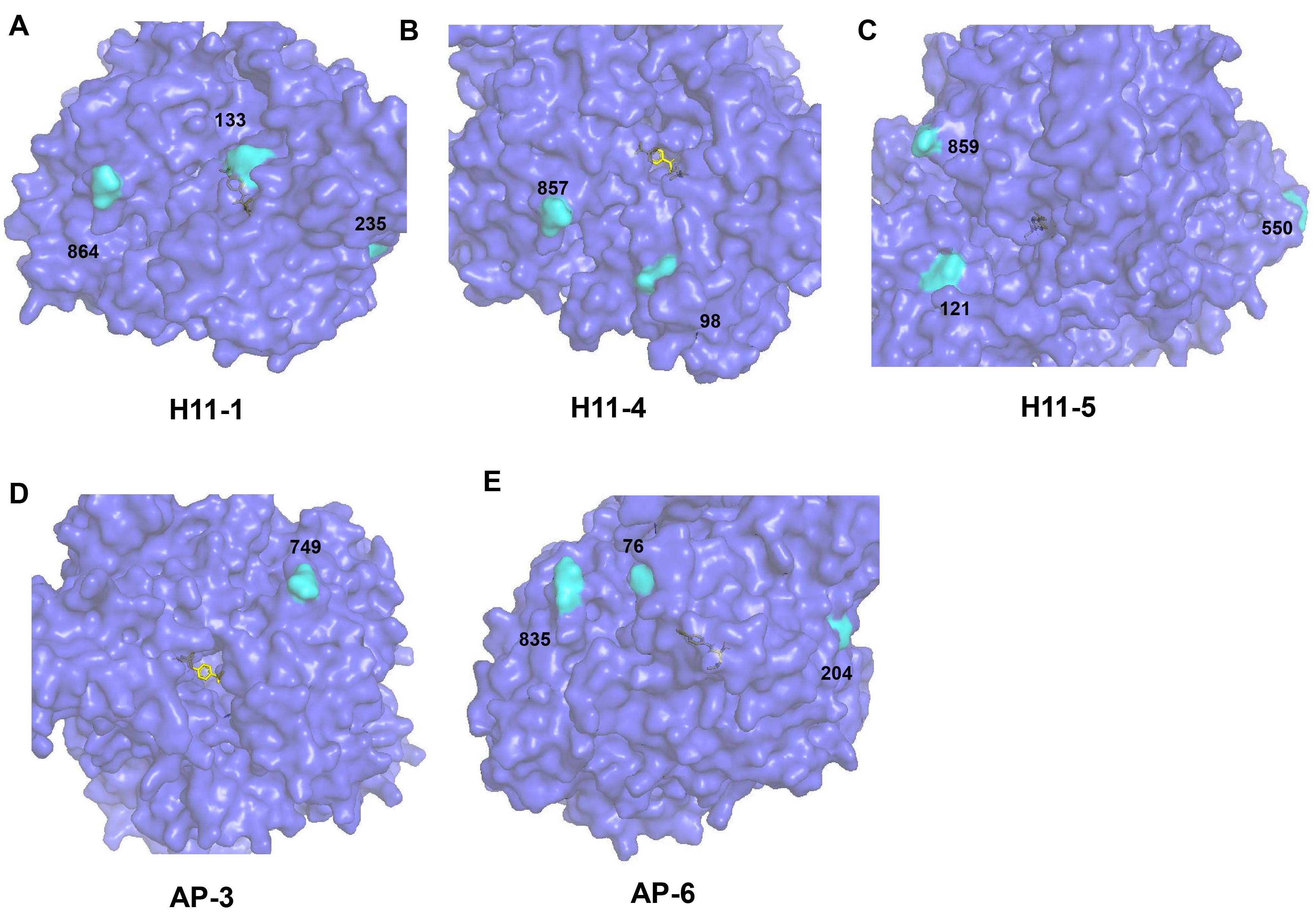
